# MindSight: A Bio-Inspired Neural Architecture for Visual Restoration via Cortical Electrical Stimulation

**DOI:** 10.1101/2025.11.12.688136

**Authors:** Yongjie Zou, Haonan Niu, Bin Zhao, Guoliang Yi, Mengchuanzhi Yang, Jiawei Ju, Jiapeng Yin, Chengyu T. Li

**Affiliations:** Lingang Laboratory; Shanghai Center for Brain Science and Brain-Inspired Technology

**Author notes:** {, }.

## Abstract

Visual impairment is a common condition worldwide, and cortical electrical stimulation is one of the approaches to aid in visual restoration. However, existing methods suffer from limited precision, flexibility, and generalization in generating the desired visual perception. In this paper, we propose a novel deep learning-based algorithm for cortical electrical stimulation, named “MindSight,” aimed at enhancing the clarity and accuracy of induced visual perceptions. Our framework introduces three key innovations: (1) A differentiable biophysical model simulating cortical state transitions under electrical stimulation, enabling end-to-end training; (2) A dual-path training architecture combining neural decoding fidelity with phosphene simulation constraints; (3) An attention-guided background gated network for input filtration and, a multi-channel activation constraint to ensure the effectiveness of electrical stimulation. We validated our approach through novel experiments with macaque monkeys, demonstrating superior performance in visual perception tasks. These results highlight the potential of our approach in assisting individuals with visual impairments.

**Code:** https://github.com/zyj9902/MindSight

## Introduction

Visual impairment is a common condition worldwide (Stevens et al. 2013). The restoration of visual perception through cortical stimulation represents a transformative frontier in neuroprosthetics, offering hope for individuals with severe visual impairments caused by retinal degeneration or optic nerve damage. Traditional approaches, such as retinal implants (Dorn et al. 2013; Stingl et al. 2015), rely on intact neural pathways from the retina to the visual cortex, limiting their utility for patients with broader damage (Fernandez and Robles 2024). This outcome has motivated a shift toward approaches that bypass the eye entirely. In particular, visual cortex stimulation has gained interest (Lozano et al. 2020; Chen et al. 2020; Maghami et al. 2014; Roelfsema, Denys, and Klink 2018; Dobelle 2000) as it can potentially benefit a wider range of blindness causes (e.g. optic nerve damage, glaucoma) where retinal devices are ineffective. Recent advances with high-channel-count micro-electrode arrays (96–1024 channels) demonstrate that intracortical stimulation can evoke shape-like percepts in both primates and humans (Chen et al. 2020; Fernández et al. 2021). Primary visual cortex (V1) contains an ordered map of the visual field and relatively large surface area, making it an attractive target for electrode implants (Beauchamp et al. 2020; Beauchamp and Yoshor 2020; Lewis et al. 2015; Tehovnik and Slocum 2013; Chen et al. 2020) that could evoke patterned visual percepts. Although these types of visual prostheses may differ significantly in terms of the entry point into the visual system, they share the same fundamental mechanism of action: through electrical stimulation of small groups of neurons, they evoke the perception of spatially localized flashes of light, called phosphenes (Brindley and Lewin 1968; Foroushani, Pack, and Sawan 2018). Despite advancements, research lacks adaptive and dynamic strategies, requiring manual determination of critical visual features and transformation of these features into suitable electrical stimulation protocols (Chen et al. 2020). Moreover, existing automated methods suffer from limited precision, flexibility, and generalization in generating the desired complex visual perception based on actual visual stimuli (Bosking, Beauchamp, and Yoshor 2017).

This work presents a novel bio-inspired algorithm, named “**MindSight**”, for dynamically orchestrating electrode stimulation patterns in visual cortex. Our work is built on five key components: (1) A differentiable biophysical model simulating cortical state transitions under electrical stimulation, enabling end-to-end training; (2) A hierarchical training framework integrating neural decoding and phosphene simulation constraints, extending hybrid autoencoder (Granley, Relic, and Beyeler 2022); (3) Background gated network (BGN) for input filtration; (4) Multi-channel activation constraint to encourage a sufficiently large subset of electrode channels to fire concurrently, rendering the stimulation effective and consistent (Oswalt et al. 2021; Bosking et al. 2022); (5) Comprehensive validation through novel primate experiments demonstrating state-of-the-art performance.

## Related Work

In biological vision, two frequently explored directions are: (1) visual encoding mechanisms—how visual stimuli elicit neural responses; and (2) visual decoding algorithms—how to reconstruct perceived visual scenes from neural signals. Extensive research has been conducted on the neural encoding mechanisms and response characteristics to visual stimuli in rodents (Turishcheva et al. 2024b,a; Li et al. 2023; Xu et al. 2023; Sinz et al. 2018) and primates (Cadena et al. 2019; Hatanaka et al. 2022; Farzmahdi, Kohn, and Coen-Cagli 2025; Shipp 2024), leading to significant advances and a clearer understanding of these processes. Similarly, substantial progress has also been made in the area of visual decoding. Pixel-level reconstructions of images from brain signals demonstrate how encoding and decoding approaches inform each other (Zhang et al. 2020, Zhang et al. 2022). Meanwhile, other studies focus on decoding semantic information (Chen et al. 2023; Ozcelik et al. 2022; Gaziv et al. 2022).

Given that the aforementioned studies primarily focus on the relationship between visual input and neural activity, either predicting natural neural responses based on given visual stimuli and brain states, or conversely decoding perceived visual input from neural activity. However, they differ substantially from the goal of visual prosthetics: determining how to appropriately intervene in the visual system to evoke desired percepts. Nonetheless, the mechanisms of visual encoding and decoding still provide important biological insights into how visual perception might be restored.

Early visual prosthetics focused on retinal stimulation, exemplified by systems like Argus II, which translates camera-captured images into electrical signals for retinal neurons (Luo and Da Cruz 2016). However, such devices require intact retinal circuitry, rendering them ineffective for patients with optic nerve damage or advanced degenerative diseases. Cortical visual prosthetics bypass these limitations by directly stimulating visual cortex neurons but initially yielded only coarse visual perceptions (Brindley and Lewin 1968; Dobelle, Mladejovsky, and Girvin 1974). There is a need to explore more complex pattern presentation (Dobelle and Mladejovsky 1974; Bosking, Beauchamp, and Yoshor 2017). Recent cortical prosthetic approaches advanced visual perception by either sequentially activating electrodes to mimic natural motion-sensitive processing (Beauchamp et al. 2020), enabling accurate letter recognition in humans, or employing high-channel-count simultaneous stimulation to evoke shape perception in monkeys (Chen et al. 2020). Meanwhile, some deep learning techniques have also been introduced into this field. A deep autoencoder-based architecture that includes a highly adjustable prosthetic vision simulation module attempts to automatically find a task-specific stimulation protocol (van Steveninck et al. 2022). Hybrid neural autoencoder (Granley, Relic, and Beyeler 2022) combines biological constraints with deep learning to generate stimulation that preserve topological relationships in V1. Another ML-based paradigm is to incorporate models of the visual system’s response into the stimulus optimization process (Grani et al. 2022).

However, these methods either remain at the theoretical simulation stage without sufficient animal experiments to validate their effectiveness, or the visual percepts elicited by the methods are imprecise, inflexible, and unclear. To address these issues, this work proposes MindSight.

## Methodology

### Overview

As shown in Figure 1, MindSight consists of three major stages. The first stage is the MUA-Image Decoder (MID), which decodes neural activity in the form of multi-unit activity (MUA) into corresponding visual images. The second stage is the Electrical Stimulation Generator (ESG), which optimizes the stimulation parameters to generate the visual perceptions as close as possible to the desired perceptions. This stage involves training through two paths of losses. The third stage, the Background Gated Network (BGN), filters out irrelevant visual stimuli, ensuring that electrical stimulation is applied only to stimuli that contain essential visual information. The biologically inspired designs of the individual modules, along with the rationale behind each, will be detailed in the corresponding subsections that follow. To provide an overview of how the algorithm operates as a whole, the high-level algorithmic workflow of the entire framework is presented in **Algorithm 1** in the **Appendix**.

**Figure 1.**
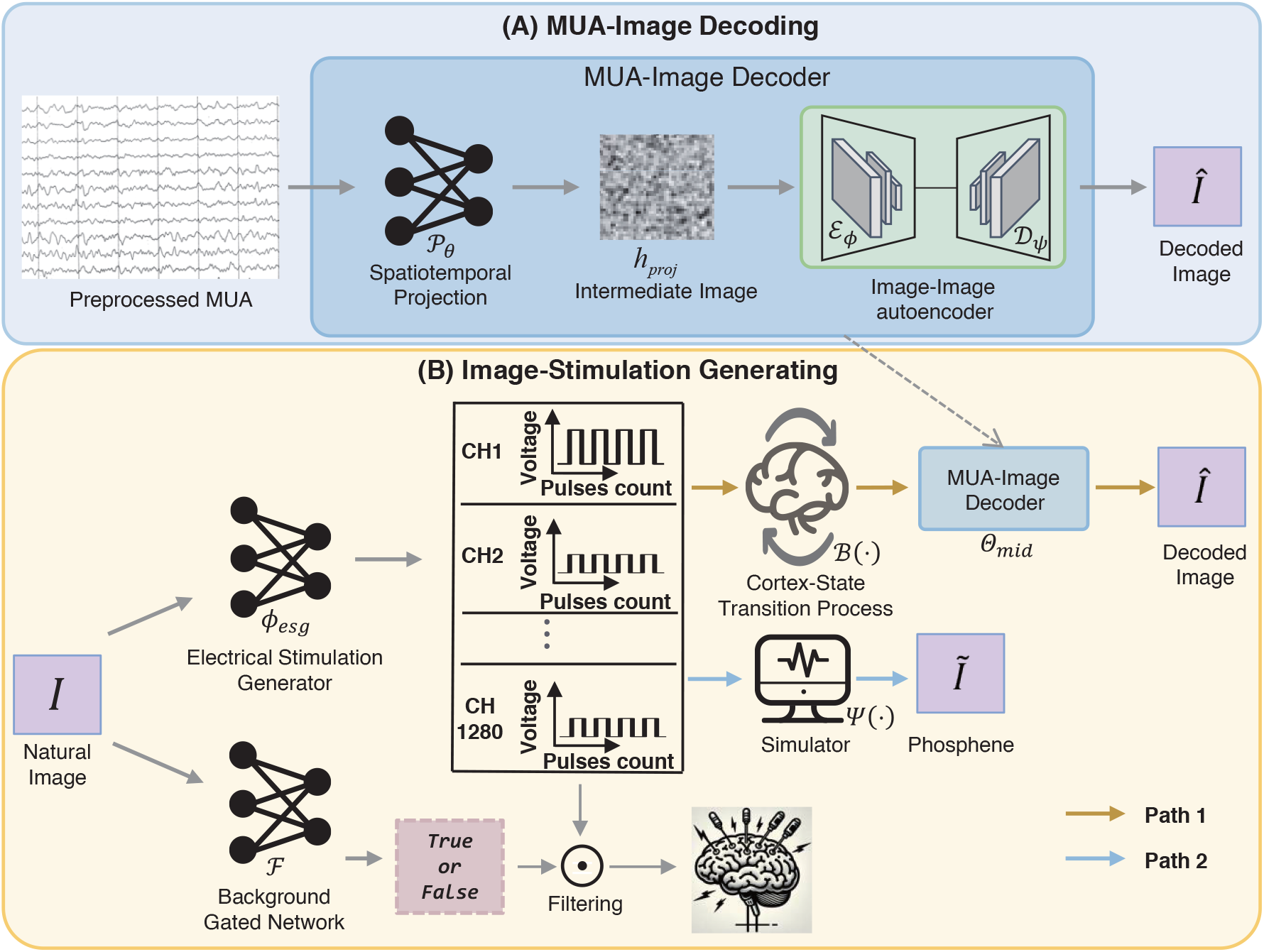
MindSight. Stage A (upper part): In MUA-Image Decoding, MUA-Image Decoder is trained to reconstruct visual stimuli from the multi-unit activity (MUA) recorded in the primary visual cortex, and the well-trained MUA-Image Decoder will be used in stage B. Stage B (lower part): In Image-Stimulation Generating, electrical stimulation generator (ESG) is responsible for producing voltage amplitudes on each electrode channel, given a real input image, and its parameters are trained via two paths. Background gated network (BGN) is trained to filter the external visual inputs, ensuring that only images containing key information will trigger cortical electrical stimulation.

### Stage A: MUA-Image Decoder (MID)

In previous work, attempts were made to decode pixel-level images from retinal ganglion cells spikes (Zhang et al. 2020) and two-photon calcium signals of the Macaque Visual Cortex (Zhang et al. 2022). However, these models do not incorporate biological visual mechanisms. To reflect the selective tuning of visual cortical neurons to different visual attributes, the modulation of receptive fields, and the feedback and integration across visual cortical areas, MUA-Image Decoder (see the upper part of Figure 1) introduces three key advancements over previous neural decoders: (1) Spatiotemporal fusion of multi-scale MUA dynamics, (2) Hierarchical attention-guided feature refinement, and (3) Biologically-plausible feature recombination through adaptive skip connections. Our architecture incorporates the Convolutional Block Attention Module (CBAM) (Woo et al. 2018) for cross-dimensional feature refinement. As shown in Figure 1, the overall calculation process of MID is as follows:

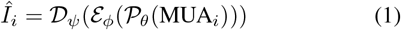

Here, *Î*_*i*_ denotes the image decoded from MUA (1280 channels × 50 ms), 𝒫_*θ*_ represents the projection network, and ℰ_*ϕ*_ and 𝒟_*ψ*_ denote the encoder and decoder of the Image-Image autoencoder, respectively. Refer to **Appendix A** for details.

### Stage B: Electrical Stimulation Generator (ESG)

The ESG (see the lower part of Figure 1) is responsible for producing voltage amplitudes on each electrode channel, given a real input image. The ESG parameters would be optimized via two paths (referred to as *Path 1* and *Path 2*) and one channel-activation constraint term, each contributing to the overall loss function:

#### 1. Path 1: MID Decoding Constraint

The purpose of this constrained pathway is to ensure that the neural state changes and responses induced by cortical electrical stimulation can be decoded—via the MID trained in stage A—into the intended visual input. This provides a theoretical guarantee of stimulation effectiveness from the perspective of biological vision systems. Otherwise, how can we ensure that the visual percepts generated by electrical stimulation are meaningfully related to the desired visual inputs?

#### 2. Path 2: Phosphene Simulation Constraint

The inclusion of this constrained pathway is also motivated by the need to satisfy established biological priors— namely, the orderly mapping between the visual cortex and phosphene perception. Without this, even at the coarse level of phosphene perception, the correctness of electrical stimulation cannot be ensured.

#### 3. Channel Activation Constraint

To ensure that sufficiently many channels within the same electrode array are activated in unison. Without this, stimulating too few electrode channels in a local region may render the stimulation ineffective—that is, no percept may be elicited in the corresponding area of the visual field.

##### Architecture of ESG

To better enable the ESG to learn the relationship between the target visual percept and the corresponding electrical stimulation parameters, it is essential to extract features from different regions of the visual target space and across multiple levels of granularity. At the same time, the design must account for the deployment and computational cost of ESG.

Depthwise-separable convolutions are employed to reduce the siz of ESG, aiming to minimize power consumption, as ESG will be integrated into the chip within the visual prosthesis system. Hierarchical processing with depthwise-separable convolutions and attention:

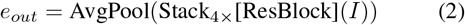

Use ℛ to explicitly denote each ResBlock, as shown below:

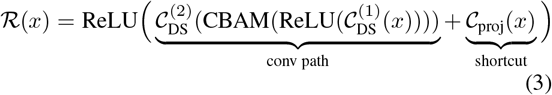

where 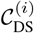 denotes depthwise-separable conv with BN.

After adaptive global pooling, we apply two fully connected layers (with LayerNorm and GELU) to predict stimulation parameters:

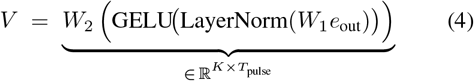

Here *W*_1_ and *W*_2_ are linear transformations mapping and *V* represents the voltage amplitudes for the *K* electrode channels. *T*_pulse_ denotes the number of pulse for each channel.

#### Path 1: MID Decoding Constraint

As shown in Figure 1, in path 1, we introduce the cortex-state transition process and the previously trained MID to guide ESG: 1. A real input image *I*_*i*_ is fed into the ESG, denoted by *ϕ*_*esg*_(·), yielding a set of voltage amplitudes across all electrode channels. 2. A cortex-state transition process model simulates how these voltage amplitudes (converted to currents) affect cortical neural states over time, producing a simulated neural response. 3. The simulated neural response is passed through the fixed-parameter MID, which reconstructs an image *Î*_*i*_.

Let Θ_*mid*_ be the fixed parameters of MID. Let ℬ (·) denote the *cortex-state transition process*, modeling the spatiotemporal changes in mua triggered by electrical stimulation. Path 1 can be summarized as:

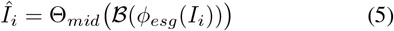

#### Cortex-State Transition Process

Suppose each of the *K* electrodes is located at spatial coordinates (*x*_*k*_, *y*_*k*_, *z*_*k*_), and let 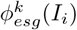 be the generated voltage amplitude on electrode *k*. If *R*_*k*_ is the impedance coefficient of channel *k*, then its injected current is approximately

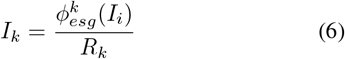

We adopt Spatiotemporal Gaussian distribution to describe how stimulation at (*x*_*k*_, *y*_*k*_, *z*_*k*_) and current *I*_*k*_ diffuses through cortical tissue over time. In the *x, y, z* directions, we assume that the diffusion rates *α* are the same, i.e., 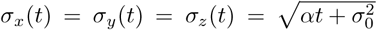 Specifically, after calculation and simplification, the electrical stimulation for channel *k* ultimately generated the following distribution:

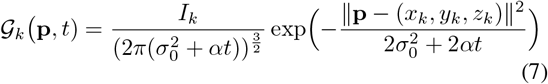

where **p** denotes a spatial coordinate, *t* represents time, *σ*_0_ is the initial standard deviation, and *α* is the diffusion rate. Summing contributions from all *K* electrodes yields

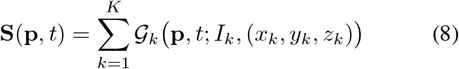

thus forming a spatiotemporal distribution of the electric field. By sampling **S**(**p**, *t*) at the corresponding recording channels over a certain time window, we obtain the simulated neural response, which is then fed into the MID. Note that *T*_pulse_ is a critical hyperparameter and Equations (7) and (8) only describes the case where *T*_pulse_ is 1. If *T*_pulse_ is greater than 1, the complex effects introduced by different pulses must also be considered. In our experiments, we set *T*_pulse_ to 1. If you find that a small number of pulses cannot activate the channel, then *T*_pulse_ needs to be increased.

#### Loss Function for Path 1

The goal of path 1 is to ensure that the images reconstructed from simulated neural responses are highly consistent with the real input images, thereby ensuring that the electrical stimulation parameters generated by ESG can induce the desired visual perception. Using the mean squared error at the pixel level, we define

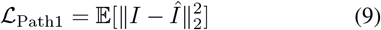

#### Path 2: Phosphene Simulation Constraint

In path 2, the same generator *ϕ*_*esg*_(·) is used, but the forward pass relies on a *phosphene simulator* (van der Grinten et al. 2024) (see the lower part of Figure 1), denoted by Ψ(·), and Ψ(·) is not trainable. The pipeline is: 1. A real image *I*_*i*_ is given to the generator *ϕ*_*esg*_(·), producing voltage amplitudes (converted to currents). 2. Generated voltage amplitudes 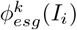, impedance coefficient *R*_*k*_ and the center positions of the receptive fields 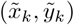 for all channels are fed into the phosphene simulator Ψ(·), which outputs a simulated phosphene image *Ĩ*_*i*_. 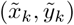 can be obtained through Receptive Field Mapping (see Section **Receptive Field Mapping** for details). Mathematically, we have

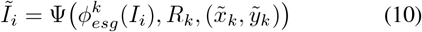

For each real input *I*_*i*_, we generate the corresponding binarized target *İ*_*i*_ (see **Appendix D: Method for Constructing Binarized Targets** for more details). The path 2 loss is then:

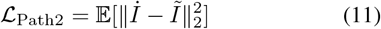

which penalizes discrepancies between the simulated and target phosphene images to adhere to the orderly mapping between the visual cortex and phosphene perception.

#### Channel Activation Constraint

Our extensive saccade experiments (Tehovnik, Slocum, and Schiller 2003; Tehovnik et al. 2005) on monkeys have shown that, stimulating too few electrode sites within a localized area may render the stimulation ineffective, i.e., no perception was induced in the corresponding visual field region (see Section **Electrical-Stimulation-Induced Saccade** for details). To encourage a sufficiently large subset of electrode channels to fire concurrently, we introduce an additional penalty:

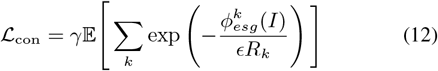

where *γ* and *ϵ* are the penalty coefficient and decay coefficient, respectively. This penalty term constrains the number of effective channels that are not stimulated to be as few as possible. In particular, when the physical locations of different channels are closer, the probability of them being stimulated simultaneously becomes higher (rather than stimulating only a subset of these channels, even though stimulating only this subset could already make ℒ_Path2_ very small).

#### Overall Loss for the ESG

By combining Path 1, Path 2, and the channel activation constraint, the parameters of ESG are trained by minimizing the total objective using Adam:

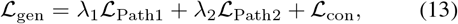

where *λ*_1_ and *λ*_2_ control relative importance of two paths.

### Stage C: Background Gated Network (BGN)

Given that multiple studies (Cogan 2008; Aungaroon et al. 2017; Larkin et al. 2022; Vatsyayan and Dayeh 2022; Vatsyayan et al. 2021) have shown excessive cortical electrical stimulation can increase the risk of neuronal necrosis, tissue damage, electrode failure, and even trigger after-discharges or seizures, we further train a BGN (see the lower part of Figure 1) to filter visual inputs. This ensures that only images containing key information trigger cortical stimulation.

BGN consists of a ResNet backbone modified with CBAM layers to dynamically refine feature maps. Given *I*_*i*_, BGN indicates whether the image contains key information:

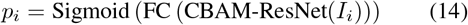

## Experiment

### Datasets

In our study, we used two self-collected datasets: **Monkey Neural-Image Viewing Dataset (NIVD)** and **Monkey Vehicle Obstacle Avoidance for Food — Neuro-Visual Dataset (MVOAF-NVD)**. Refer to **Appendix C** for more description of these two datasets. For details on how NIVD and MVOAF-NVD were collected, refer to **Appendix C.1, Appendix C.2**, and Supplementary video 1.

### Implementation Details

For all hyperparameter choices and implementation details related to data preprocessing and the training of MindSight, please refer to **Appendix B**.

### Receptive Field Mapping

We used a fast RF mapping method similar with (Fiorani et al. 2014) to get the receptive fields of neurons. Refer to **Appendix E** for more details.

### Electrical-Stimulation-Induced Saccade

We followed previously reported methods (Chen et al. 2020) to conduct electrical-stimulation-induced saccade experiments on our monkeys. We found that at least 16 channels per electrode are required to elicit effective phosphene perception, which motivated the design of Channel Activation Constraint. For more saccade experiments details, refer to **Appendix F** and the corresponding Table S1.

### Behavioral Validation Experiments in Monkeys

To validate the efficacy of MindSight, we designed a delayed match-to-sample (DMS) task and a driving-based obstacle-avoidance foraging (DOAF) task. Refer to **Appendix G** and Figure S1 for more description of these two tasks.

## Results

For MindSight, it is mainly necessary to validate the results of the following modules it contains: 1. Decoding performance of MID, as it indirectly affects the performance of ESG; 2. Phosphenes generated by ESG; 3. Performance of ESG in DMS; 4. Performance of ESG and BGN in DOAF.

### Performance of MID

We evaluated the performance of MID on the test set of the **NIVD** (4,037 cropped images with corresponding MUAs). Figure 2a demonstrates MID’s decoding results on some examples. As shown in Figure 2b, MID was compared with several state-of-the-art pixel-level decoding models—SID (Zhang et al. 2020), UNet-based model (Zhang et al. 2022), and DGMM (Du et al. 2018)—using evaluation metrics including mean square error (MSE) that describes the absolute difference of every pixel, the structural similarity index measure (SSIM) that captures the details or image distortion (Wang et al. 2004), and the peak-signal-to-noise-ratio (PSNR) that characterizes the global quality. For clarity, we utilized normalized model performance. Specifically, each model’s score for every test sample was normalized against MID’s scores, where the ratio for MID is defined as 1. Values above 1 indicate better performance than MID, while values below 1 indicate inferior performance. For the other three models, we calculated the mean and variance of their normalized performance across the entire test set. Compared to the other three models, MID from MindSight exhibited superior performance across all three metrics.

**Figure 2.**
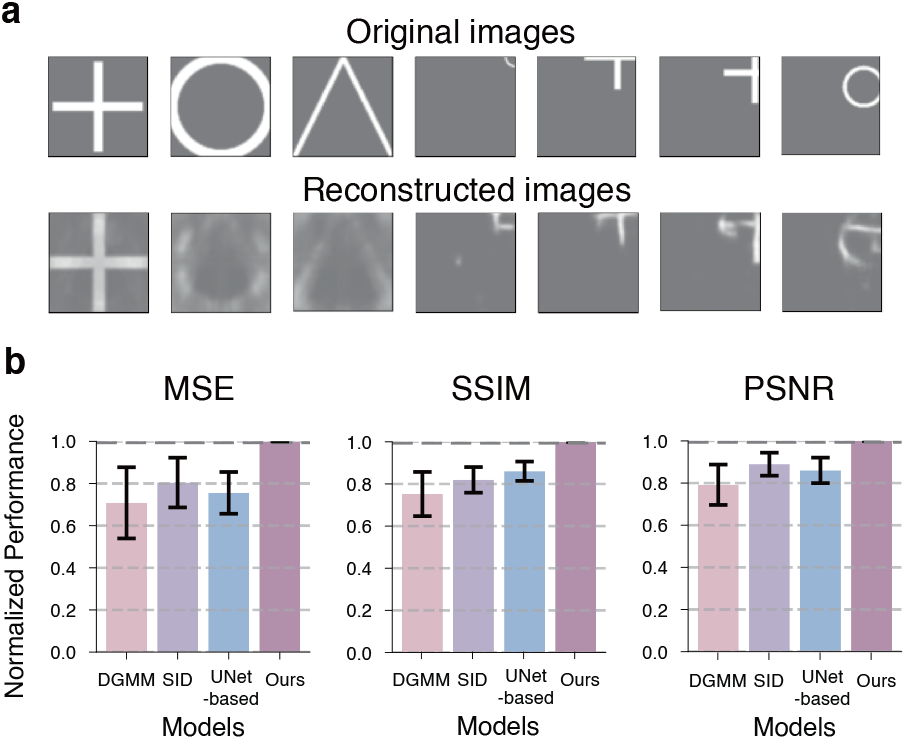
Decoding Performance Comparisons on NIVD Test Set. a. Some examples of decoded images. **b**. We evaluated four models (ours, DGMM, SID, and UNet-based) on the NIVD test set using three evaluation metrics: MSE, SSIM, and PSNR. Error bars represent standard deviation of the mean normalized performance.

### Phosphenes Evoked by ESG

We also evaluated the simulated phosphenes evoked by ESG in both DMS and DOAF. As shown in Figure 3, we assessed the theoretical phosphenes that would be induced by the stimulation parameters generated by ESG when aiming to evoke different desired visual percepts in DMS for VM15, VM22, and VM23. The phosphenes evoked by ESG vary across monkeys (Table 1 provides a quantitative summary for each monkey), due to differences in electrode implantation sites, which in turn lead to variations in receptive fields (see **Appendix E** for more details). These differences in electrode and receptive field distributions influence the training of MindSight, as described in Equations (7), (8), and (10). The results for VM15 are significantly worse than those for VM22 and VM23, primarily due to the suboptimal distribution of electrode implantation sites. The phosphenes evoked by ESG in DOAF will be detailed in Section **Performance of ESG and BGN in DOAF**.

**Table 1:** Quantitative Results for Figure 3. For each monkey, we compute the similarity between each desired visual percept and the corresponding phosphene evoked by ESG. Since the focus here is primarily on shape and contour, the original visual percept images are first preprocessed (e.g., binarization) before computing the Intersection over Union (IoU) and Dice Similarity Coefficient with the phosphenes.

| Visual percepts | VM22 |  | VM23 |  | VM15 |  |
| --- | --- | --- | --- | --- | --- | --- |
|  | IoU | Dice | IoU | Dice | IoU | Dice |
| Triangle | 0.60 | 0.75 | 0.57 | 0.73 | 0.43 | 0.60 |
| Cross | 0.64 | 0.78 | 0.68 | 0.81 | 0.48 | 0.65 |
| Circle | 0.61 | 0.76 | 0.57 | 0.73 | 0.48 | 0.64 |
| Rectangle | 0.52 | 0.68 | 0.46 | 0.63 | 0.36 | 0.53 |
| bar (0°) | 0.76 | 0.86 | 0.64 | 0.78 | 0.61 | 0.76 |
| bar (45°) | 0.69 | 0.82 | 0.73 | 0.84 | 0.58 | 0.74 |
| bar (135°) | 0.67 | 0.80 | 0.70 | 0.82 | 0.47 | 0.64 |
| bar (90°) | 0.67 | 0.80 | 0.60 | 0.75 | 0.49 | 0.66 |

**Figure 3.**
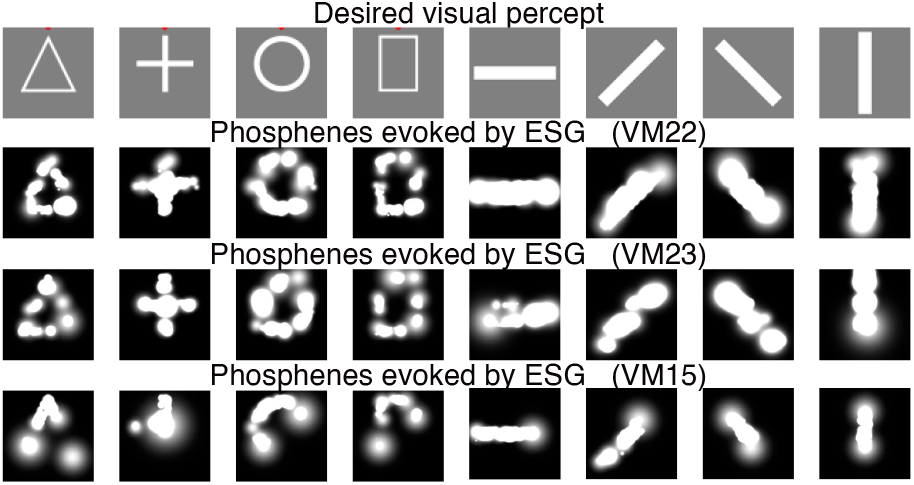
Phosphenes Evoked by ESG in DMS. The first row shows eight possible visual percepts in DMS. The second, third, and fourth rows show the phosphenes that ESG would theoretically evoke for VM22, VM23, and VM15, re-spectively, when targeting each desired visual percept.

### Performance of ESG in DMS

The ESG trained on NIVD was first validated in DMS (as outlined in Section **Behavioral Validation Experiments in Monkeys**). We evaluated model efficacy by measuring the monkey’s behavioral success rate in DMS, where the choice relies entirely on electrical stimulation, with no visual cues provided. Figure S1 displays four-class DMS, with the two-class DMS differing only in the final selection step between two shapes instead of four shapes. Figure 5a shows the dynamic performance of behavioral success rates in a two-class DMS during one session. With electrical stimulation from the ESG, the monkey maintained high accuracy (red region). However, when the stimulation was removed, the success rate immediately dropped to chance level (green region). As shown in Figure 5b, the results of MindSight’s ESG in two-class DMS were compared with those from another Chen’s study (Chen et al. 2020). Both experiments followed nearly identical protocols, except that Chen used some different shapes, such as “T” and “L”. We validated our method using two monkeys (monkeys 22 and 23), while Chen used monkeys A and L, with results statistically aggregated over 10 sessions per monkey. The comparison reveals that our method achieves higher stability. Moreover, compared to (Chen et al. 2020), MindSight also offers greater flexibility, as it can directly adapt to arbitrary inputs without the need for additional manual intervention. Such robustness and flexibility are crucial for the future application of visual prosthetics. We additionally demonstrated the monkeys’ performance in four-class DMS (which has not been explored in (Chen et al. 2020)). Figure 5c illustrates the dynamic performance of behavioral success rates in one session for monkeys 22 and 23, respectively. Similarly, a control group without electrical stimulation was set up, confirming that MindSight’s correct rate far exceeded the chance level (25%). Supplementary videos 2 and 3 demonstrate examples of MindSight applied to 2-class (Monkey 22) and 4-class (Monkey 23) DMS, respectively, showing the monkeys performing continuous correct trials. See **Appendix H** for details of Supplementary Videos 2–3.

### Performance of ESG and BGN in DOAF

The ESG (trained on MVOAF-NVD) was validated in DOAF (as outlined in Section **Behavioral Validation Experiments in Monkeys**). Because a full session is lengthy, Supplementary Video 4 presents only segments highlighting MindSight’s performance at key moments (e.g., when the monkey observes markers on walls, food dispensers, or irrelevant background). In the video, left panel shows head-mounted camera footage (on a head-fixed monkey), with the red dashed box representing a 64×64 region corresponding in both size and position to the receptive field covered by implanted electrodes. Upper-right display shows the content within this red dashed box, which serves as the input to ESG/BGN. Lower-right panel simulates the visual perception evoked by the electrical stimulation commands generated by ESG (Equation (10)), combined with BGN outputs. For easier viewing, Figure 4 presents some frames from Supplementary Video 4 as illustrative examples. BGN-triggered stimulation activates only during key cues (e.g., markers, obstacles, and food dispensers), filtering out irrelevant information to avoid the risks associated with excessive electrical stimulation. Notably, despite variations in the shape, angle, and position of markers and food dispensers as the cart moves, MindSight consistently evoked effective visual perceptions.

**Figure 4.**
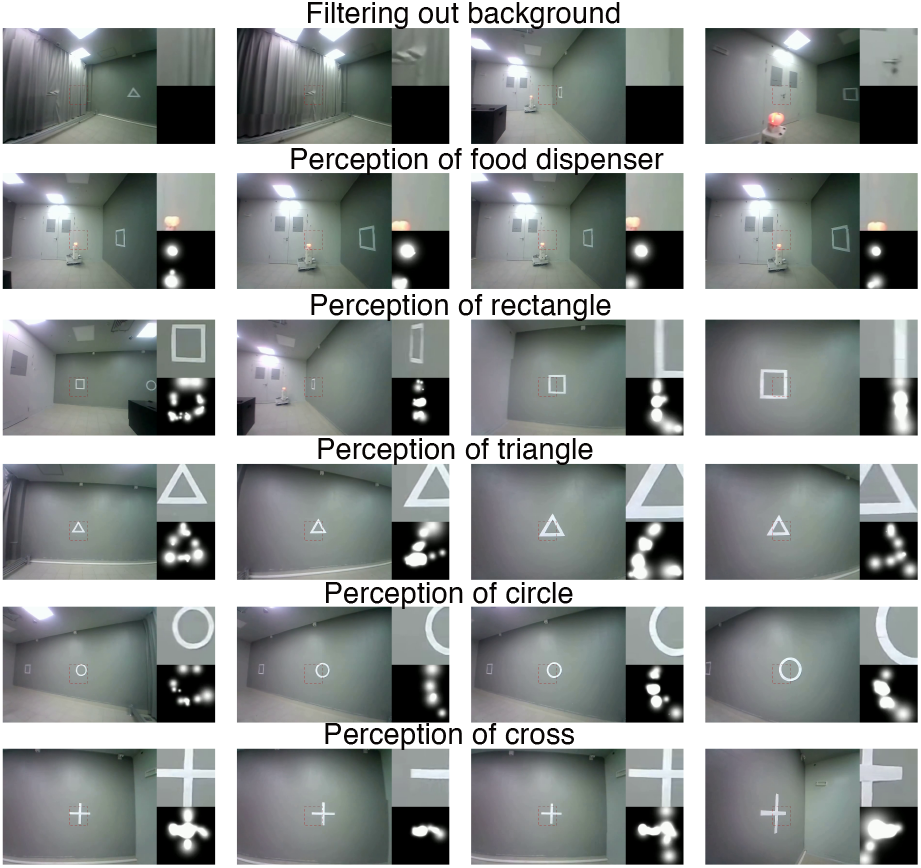
Examples of Phosphenes Evoked by ESG in DOAF. The first row shows the results of background filtering, where irrelevant information such as door handles, curtains, and walls does not trigger electrical stimulation. Rows two through six illustrate the stimulation evoked for the food dispenser and various markers at different angles and distances as the cart moves.

**Figure 5.**
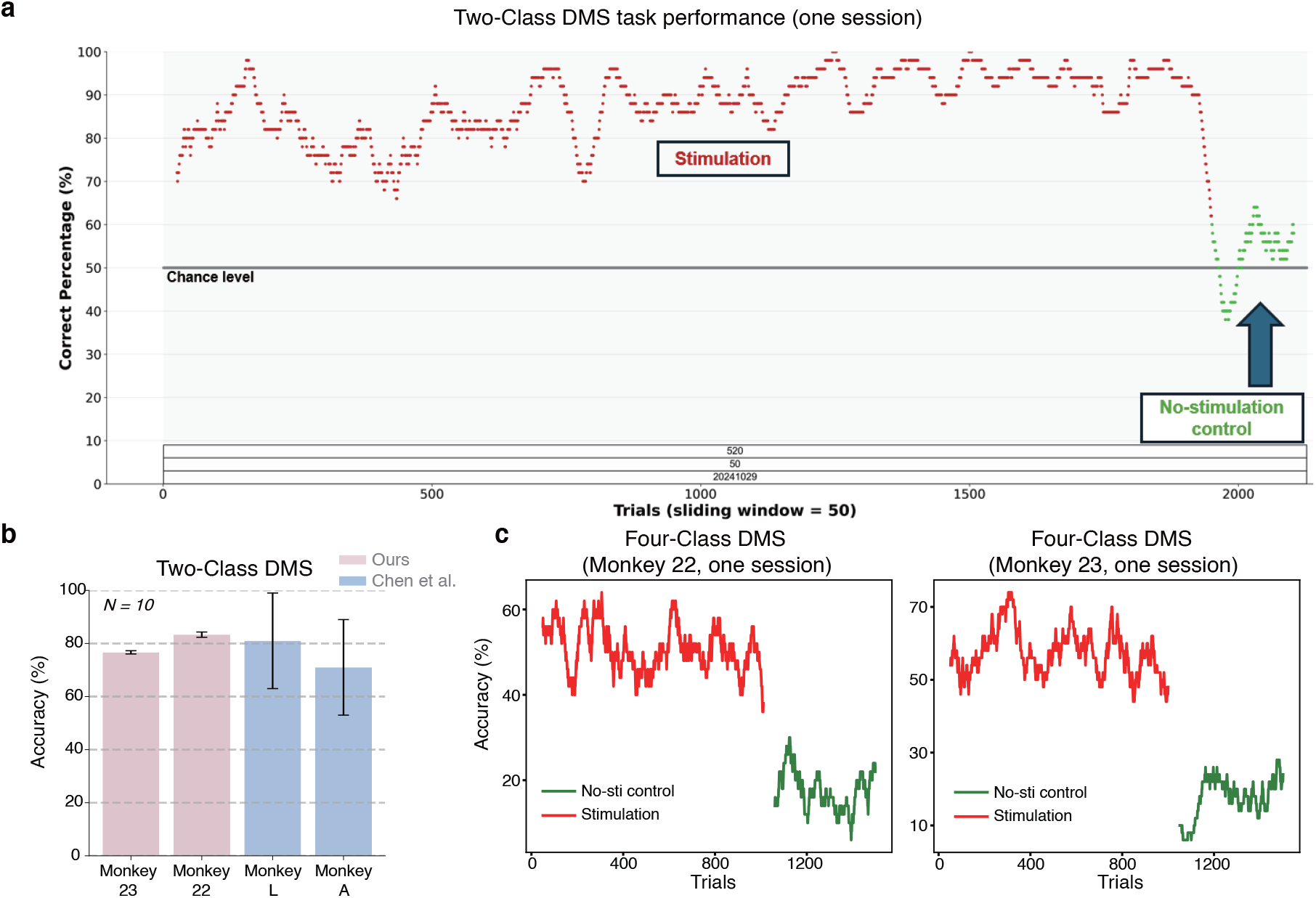
Validation on the DMS Task. a. Dynamic performance of MindSight in two-class DMS task during one session. **b**. Performance comparisons in two-class DMS task. Error bars represent the SEM (Standard Error of the Mean). N, number of sessions for each monkey and model. **c**. Dynamic performance of MindSight in four-class DMS task during one session.

## Conclusion

We present MindSight, a framework for visual restoration via cortical electrical stimulation. By integrating differentiable biophysical modeling with dual-path training, Mind-Sight outperforms other approaches and shows good performance with comprehensive validation through novel primate experiments. For additional discussion, see **Appendix I**.

## Acknowledgments

This work was supported by National Science and Technology Innovation 2030 Major program (No. 2021ZD0203601), National Natural Science Foundation of China (No. 32221003, No. 32221003), Shanghai Municipal Science and Technology Major Project (No. 2018SHZDZX05, No. 2021SHZDZX), National Key R&D Program Key Scientific Issues of Transformational Technology (No. 2019YFA0709504), Shanghai Pilot Program for Basic Research-Chinese Academy of Science, Shanghai Branch (No. JCYJ-SHFY2022-010), Lingang Laboratory (No. LG202105-01-01, No. LG202105-01-11, No. LG-GG-202402-06, No. LGL-5925), Shanghai Sailing Program (NO. 22YF1460900, No. 23YF1461000), Shanghai Post-doctoral Excellence Program (No. 2023021), and Shanghai Yang Fan Foundation (No. 24YF2730700).

## Appendix

## A Detailed Architecture of MUA-Image Decoder (MID)

As shown in Figure 1 (in the main text), the overall calculation process of MID is as follows:

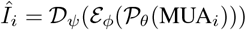

Here, *Î*_*i*_ denotes the image decoded from MUA (1280 channels × 50 ms), 𝒫_*θ*_ represents the projection network, and ℰ_*ϕ*_ and 𝒟_*ψ*_ denote the encoder and decoder of the Image-Image autoencoder, respectively.

Specifically, MUA is first processed through a spatiotemporal projection (𝒫_*θ*_). Each channel’s waveform is processed using a one-dimensional convolution layer, followed by batch normalization (Ioffe and Szegedy 2015) and adaptive global temporal pooling to obtain the feature vector *h*. This feature is then passed through a fully connected layer and reshaped into an intermediate image *h*_proj_. The detailed computation process is as follows:

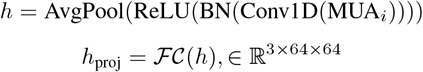

The intermediate image representation is then fed into an autoencoder (Attention-Enhanced Encoder ℰ_*ϕ*_ combined with Biologically-Inspired Decoder 𝒟_*ψ*_) to reconstruct an image. The Attention-Enhanced Encoder performs four-stage feature extraction through an EncoderBlock composed of Conv2D → CBAM → MaxPool, capturing multi-scale and hierarchical refined features *e*_*j*_ from the intermediate image, where *j* represents the layer number of the feature. The detailed computation process is as follows:

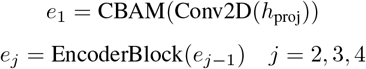

To capture the selective tuning of visual cortical neurons to different visual attributes and the modulation of receptive fields, the Biologically-Inspired Decoder employs spatiotemporal attention to associate distinct parts and properties of the visual input with neural spiking activity across different channels and time points. To mimic the feedback and integration of information across visual cortical areas, the decoder further recombines multi-scale features through channel-wise concatenation with skip connections, enabling spatiotemporal fusion of multi-scale MUA dynamics. The detailed computation process is as follows:

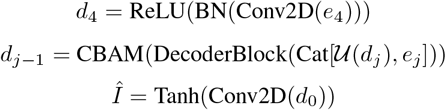

Here, 𝒰 consists of bilinear upsampling and Conv2D; DecoderBlock denotes (Conv2D→BN→ReLU)*2; *e*_*j*_ denotes the output of ℰ_*ϕ*_, which encodes multi-scale features of the intermediate image; Cat represents channel-wise concatenation with skip connections; and *j* = 4, …, 1 indicates the feature level.

Mean squared error was calculated between output decoded images *Î*and the original images *I* to train MID.

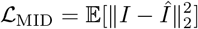

## B Implementation Details

Electrophysiological data is processed using the SpikeInter-face and Mountainsort-5 toolbox (Chung et al. 2017; Buc-cino et al. 2020) to extract multi-unit activity (MUA) from the white-band signal. Specifically, the threshold for spike detection when applied to whitened data is 4.3 standard deviations. Subsequently, the MUA is downsampled from a 20kHz sampling frequency to 1kHz, and the 1280 channels with the highest firing rates are selected. Each 64×64 image corresponds to an MUA shape of 50×1280, as the MUA is taken over a 50ms period. We chose a 50ms MUA window as the decoding input for two main reasons. First, primary visual cortex exhibit initial response windows of 20– 100ms, with neurons encoding maximal information within tens of milliseconds. Spike/PSTH counts show peak information entropy between 30–70ms. Second, guided by this biological basis, we narrowed the hyperparameter search range and ultimately identified 50ms as the optimal value. This data is used to train the MID, with a batch size of 32, a learning rate of 0.001, and the Adam optimizer to find the best model on the validation set over 1000 epochs, followed by testing on the test set.

### Algorithm 1: Algorithmic workflow of the entire framework

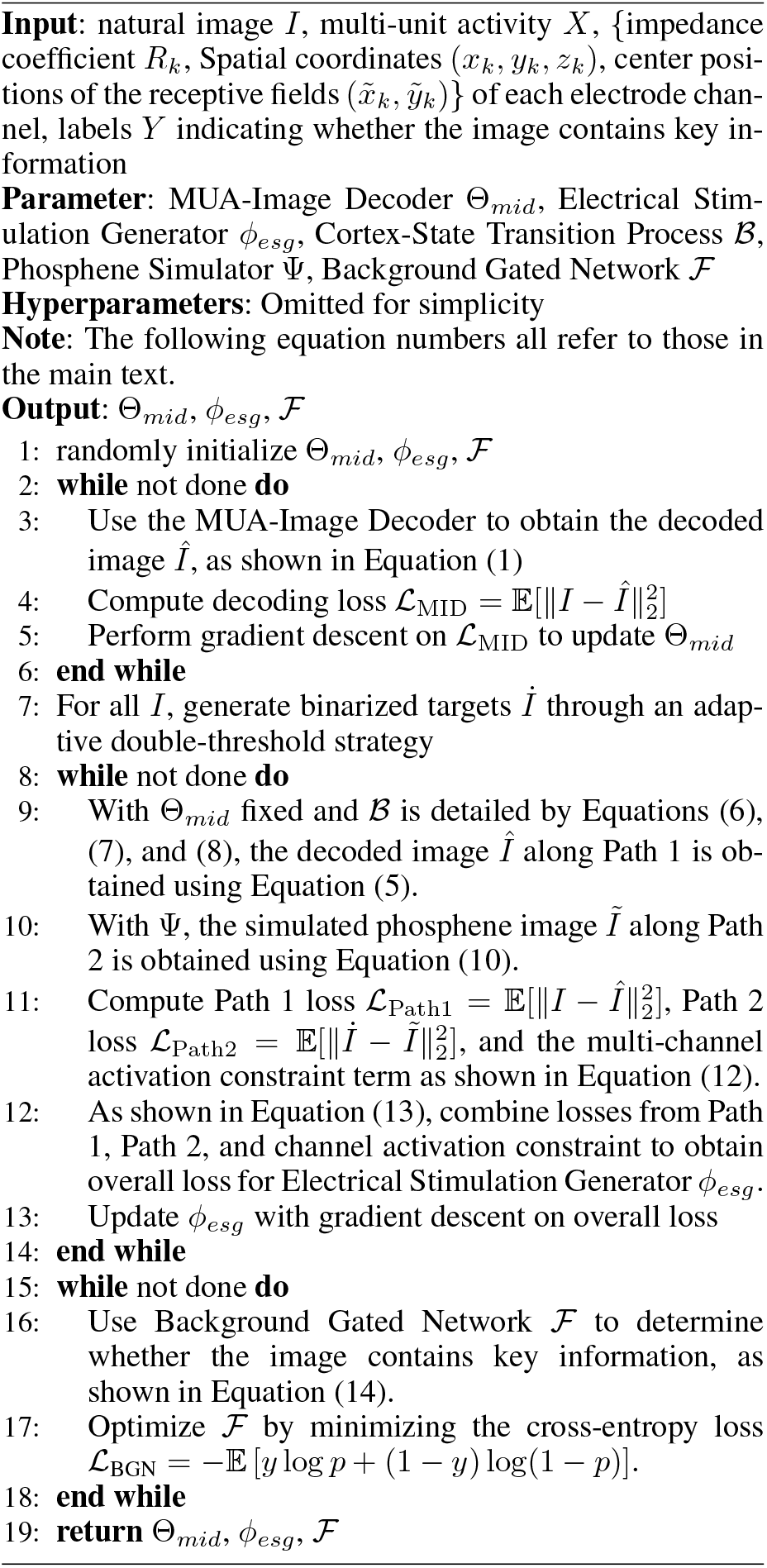

The model parameters from the trained MID are then used to train the ESG, which involves numerous hyperparameters. Based on extensive impedance testing and statistical analysis across electrodes, we set the impedance coefficient *R*_*k*_ for each electrode channel to 10000. However, In practice, stimulation-induced impedance variations (influenced by factors such as electrode materials, tissue properties, and implantation duration) exhibit unpredictable complexity, currently excluded in modeling but prioritized for future work. During the cortex-state transition process, the number of pulses *T*_pulse_ is 1, the diffusion rate *α* is 2.0, and the initial standard deviation *σ*_0_ is 5.0. For setting the value of *α* and *σ*_0_, we first established a reasonable range based on (Wu et al. 2023), and then identified the optimal value through hyperparameter search. In the multi-channel activation constraint, the penalty coefficient *γ* is set to 5e-3, and the smoothing coefficient *ϵ* is set to 3e-6. For the loss summation, *λ*_1_ is 0.3 and *λ*_2_ is 0.7. During ESG training, the batch size is 32, the learning rate is 0.001, and the Adam optimizer is used to find the best model on the validation set over 1000 epochs, followed by testing on the test set.

During BGN training, the batch size is 32, the learning rate is 0.001, and the Adam optimizer is used to find the best model on the validation set over 50 epochs, followed by testing on the test set.

## C Details of the dataset

### 1. Monkey Neural-Image Viewing Dataset (NIVD)

This dataset consists of neural signals and corresponding visual stimuli recorded while three monkeys (VM15, VM22, and VM23) viewed eight different shapes. The training set, validation set, and test set account for 80%, 10%, and 10%, respectively. 2. **Monkey Vehicle Obstacle Avoidance for Food — Neuro-Visual Dataset (MVOAF-NVD)**: This dataset includes neural signals and corresponding visual stimuli recorded while three monkeys (VM10, VM14, and VM23) drove a vehicle and avoided obstacles to obtain food in a sealed room. The dataset split is the same as that of NIVD. All experimental protocols involved in this study were approved and conducted in accordance with all relevant ethical regulations.

### C.1 Details of Collecting the NIVD Dataset

The 1024-channel recording system (RHD2000, Intan Technologies) is connected to the equipment via a custom-designed adapter board. Each 128-channel headstage, containing two digital electrophysiology interface chips (Intan Technologies RHD2164, 64 channels), connects to the adapter board through a 64-pin Molex connector, with the ground wire soldered to the GND pin near index 127. We simultaneously recorded with 4 sets of devices to achieve 4096-channel recording. During the recording, the monkeys do not need to fixate on a central point (diameter 1.0°). In each trial, a shape stimulus is presented in the center of the gray screen for 1000ms. While the monkeys freely view the screen, a wearable eye-tracking system continuously captures the position of the eye’s gaze center. The inter-trial interval is 3000ms, after which a new trial begins. The eight different shape stimuli (with randomly varying sizes) appear randomly in each session’s trials. At the beginning of each trial, an event marker is sent to the recording system. The electrophysiological records and corresponding visual inputs are paired into NIVD samples through event markers and global time. Note: The visual input captures the 64 × 64-pixel region below the gaze center of the full screen at the corresponding moment (captured by the eye-tracking system). The 64 × 64 pixels are calculated based on the receptive field region covered by the implanted electrodes, screen resolution, and the distance between the monkey’s eyes and the screen, ensuring that this pixel region matches the receptive field region of the recording channels as closely as possible.

**Table S1:**
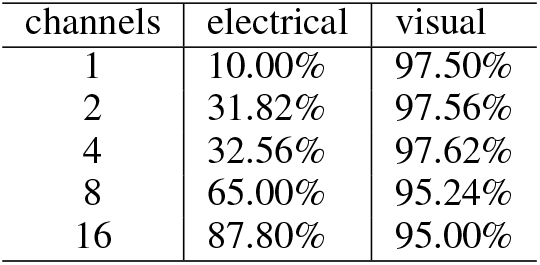
Behavioral accuracy in the saccade-to-phosphene task (first column) and the visual saccade task (second column), with each row representing a different number of channels for electrical stimulation.

### C.2 Details of Collecting the MVOAF-NVD Dataset

Supplementary video 1 (in Supplementary Material) provides an overhead perspective of a single trial in MVOAF-NVD. Using a recording system similar to NIVD, the monkey sits in a vehicle and uses a joystick to control its movement, avoiding obstacles in the room to reach the food dispenser and obtain food. Various shaped markers were attached to the walls and obstacles in the room. The visual input is obtained by a camera mounted on the monkey’s head. Similarly, electrophysiological records and corresponding visual inputs are paired into MVOAF-NVD samples through event markers and global time.

## D Method for Constructing Binarized Targets

In **Path 2: Phosphene Simulation Constraint**, for each real input *I*_*i*_, we generate the corresponding binarized target *İ*_*i*_ (with the foreground being the region that requires attention from electrical stimulation) through an adaptive double-threshold strategy: When color information is strong, it prioritizes segmentation of chromatic features to construct the binary mask; otherwise, the image is converted to grayscale, and the Otsu thresholding method (Otsu et al. 1975) is applied in combination with secondary percentile filtering.

## E Electrode Implantation and Receptive Field Mapping

We used a fast RF mapping method similar with (Fiorani et al. 2014) to get the receptive fields of neurons. Figure S2 illustrates the electrode implantation and receptive field distributions for monkeys VM22 and VM23, respectively.

## F Electrical-Stimulation-Induced Saccade

We followed previously reported methods (Chen et al. 2020) to conduct electrical-stimulation-induced saccade experiments on our monkeys. Specifically, we simultaneously stimulated varying numbers of channels on the same electrode to induce phosphenes. As shown in Table S1, by comparing task performance under visual stimulation versus electrical stimulation, we found that at least 16 channels on a single electrode must be stimulated simultaneously to achieve accuracy comparable to that under visual stimulation. This reflects the minimum number of channels required per electrode to elicit effective phosphene perception, which motivated our design of multi-channel activation constraint.

**Figure S1:**
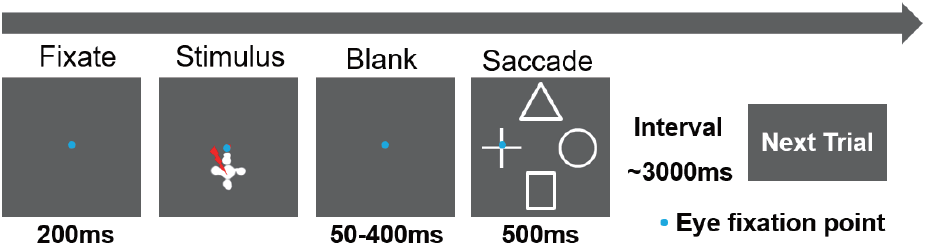
llustration of the DMS Task.

## G Task Paradigm

We designed a delayed match-to-sample (DMS) task and a driving-based obstacle-avoidance foraging (DOAF) task to validate the efficacy of MindSight.

### DMS

As illustrated in Figure S1, the monkey is first required to fixate on a central red dot (1.0°) for 200ms. Then, a randomly selected shape is fed into ESG to generate electrical stimulation parameters (executed by Intan RHS2116based device). It is important to note that during this “stimulus” phase, the screen remains blank and the monkey’s eyes are open. The “cross shape” shown under “stimulus” in Figure S1 is merely illustrative of the stimulation event and is not actually visible to the monkey. After stimulation, a 50– 400ms delay (gray screen with a fixation point, and fixation must be maintained) preceded a selection phase: saccading to the matching shape among four options and holding fixation for 500ms. Inter-trial interval: 3000ms.

### DOAF

Blindfolded monkeys navigated using a head-mounted camera feeding images to MindSight. When the BGN detected critical cues (e.g., obstacles, food dispensers), electrical stimulation triggered by the ESG enabled blind-folded monkeys to navigate obstacles and obtain food via cortical-stimulation-induced perception.

## H Details of Supplementary Videos 2 and 3

The video is divided into four panels: the larger left section displays real-time screen footage, the upper-right panel shows camera-captured scenes, the right half of the lower-right panel shows tracking the monkey’s gaze center via an eye tracker, and the left half of the lower-right panel simulates the visual perception evoked by the ESG’s electrical stimulation commands (based on Equation (18) in the main text). Note that this simulated perception is invisible to the monkey and is included solely for visualization and analysis.

## I Discussion

Performance validation in DOAF is confined solely to simulations of electrical stimulation due to the current electrode impedance issues; real-world electrical stimulation testing awaits re-implantation. Human validation remains challenging due to medical device constraints. Given the anatomical and perceptual similarities between macaques and humans, we used macaques as the subjects for this study. Our results demonstrate the potential of MindSight to propel the widespread application of visual prostheses in the future. Future work will focus on two main aspects: firstly, utilizing neural recordings obtained before and after cortical stimulation through simultaneous stimulation and recording techniques to study and optimize a more complex and realistic Cortex-State Transition Process; secondly, exploring how to dynamically adjust parameters during long-term stimulation to adapt to changes in biological states and electrode conditions.

**Figure S2:**
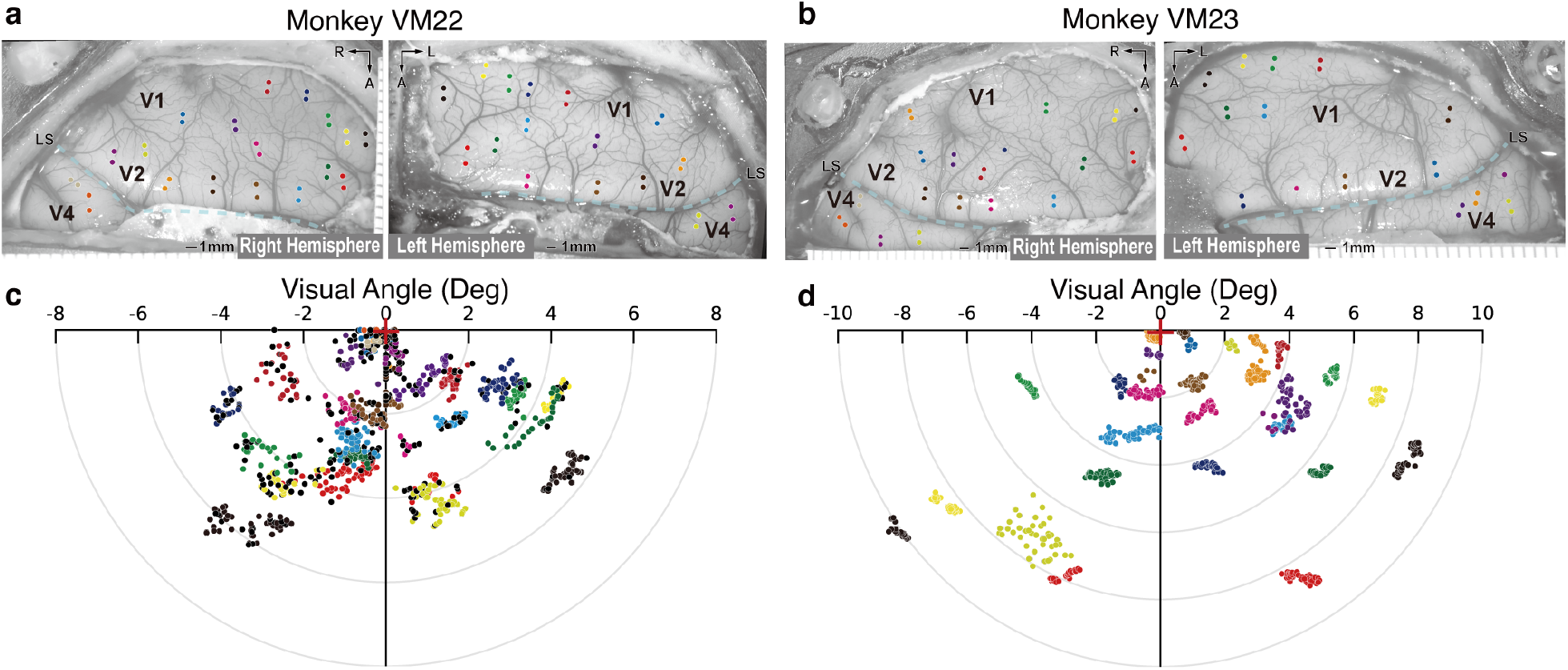
Surgery Annotation and Receptive Fields. **a** and **b** show surgery photograph of VM22 and VM23, respectively, overlaid with annotated brain regions and electrode implantation sites. Different colors represent different electrodes, and each electrode has two points because it consists of two shanks; each shank contains 64 channels. **c** and **d** show the receptive fields of VM22 and VM23, respectively (with the red cross indicating the center of each receptive fields), where each dot represents the receptive field location of a single channel from an implanted electrode (matched by color with **a** and **b**). Each electrode contains a total of 128 channels.

## J Data and Code Availability

You can find the data and code at https://github.com/zyj9902/MindSight.

